# Retinal factors of visual sensitivity in the human fovea

**DOI:** 10.1101/2021.03.15.435507

**Authors:** Niklas Domdei, Jenny L. Reiniger, Frank G. Holz, Wolf Harmening

## Abstract

Humans direct their gaze towards visual objects of interest such that the retinal images of fixated objects fall onto the fovea, a small anatomically and physiologically specialized region of the retina displaying highest visual fidelity. One striking anatomical feature of the fovea is its non-uniform cellular topography, with a steep decline of cone photoreceptor density and outer segment length with increasing distance from its center. We here assessed in how far the specific cellular organization of the foveola is reflected in visual function. Increment sensitivity to small spot visual stimuli (1 x 1 arcmin, 543 nm light) was recorded psychophysically in 4 human participants at 17 locations placed concentric within a 0.2-degree diameter around the preferred retinal locus of fixation with adaptive optics scanning laser ophthalmoscopy based microstimulation. While cone density as well as maximum outer segment length differed significantly among the four tested participants, the range of observed threshold was similar, yielding an average increment threshold of 3.3 ± 0.2 log10 photons at the cornea. Thresholds were correlated with retinal eccentricity, as well as cone density and outer segment length. Biophysical simulation allowed to develop a model of foveal sensitivity based on these parameters, explaining at least 37% of the observed threshold variability. Based on high reproducibility in replicate testing, the residual variability is assumed to be caused by individual cone and bipolar cell weighting at the specific target locations.

## Introduction

The presence of a fovea in the retina of some birds, fish, reptiles and primates like humans (Slonaker, 1897), provides an outstanding opportunity to study the linkage between a sensory system’s cellular architecture and its function. In humans, the fovea features several morphologic specializations as a result of a complex series of events that are initiated during gestation and completed in childhood (Yuodelis and Hendrickson, 1986; Zhang et al., 2020). The network of retinal blood vessels organizes itself around an avascular zone concentric within the fovea (Provis et al., 2013) and the foveal pit emerges as post-receptoral neurons migrate laterally. As a consequence, the foveola, the central 0.6 degree diameter of the fovea (Cuenca et al., 2020), is completely free of overlying neural tissue, favoring undisturbed light catch. Cone photoreceptors migrate inwards to form a lattice of tightly packed receptors at their smallest diameter found anywhere in the retina (Curcio et al., 1990), and simultaneously, the pigment-laden cone outer segments elongate (Yuodelis and Hendrickson, 1986), leading to a peak in optical density at the foveola (Elsner et al., 1993; Marcos et al., 1997). Psychophysically, photopic light sensitivity drops rapidly outside of the foveola (Choi et al., 2016).

At close inspection, the topography and density of cone photoreceptors within the foveola is highly variable between individuals (Wang et al., 2019) and it is not known, to what extent the individual cellular mosaic and the exact foveal topography is related to visual sensitivity. In patients with inherited retinal degeneration, for example, light sensitivity in the fovea was similar to a healthy control group and showed declined sensitivity only when peak cone density was reduced to less than 60% of the average (Foote et al., 2018). Sensitivity to light is assumed to be conveyed primarily by parasol ganglion cells (Takeshita et al., 2017), supported by the observation that spatial summation is correlated with the receptive field size of parasol ganglion cells (Volbrecht et al., 2000). In the foveola, however, the spatial summation area is less than 5 arcmin^2^, between the dendritic field size of parasol and midget ganglion cells (Tuten et al., 2018). Thus, a visual stimulus smaller than the spatial summation area might reveal an otherwise obscured relationship between the peaked topography of the detector array in the foveal center and its visual sensitivity.

Concomitant to the peaking spatial sampling capacity at the fovea, motor circuits of the brain stem generate eye and head movements to bring the retinal images of objects of interest detected in the periphery into the foveola (Goldberg and Walker, 2013; Poletti et al., 2013), where they fall on a distinct bouquet of only a handful cones, the preferred retinal location of fixation (PRL or “optimal locus”) (Nachmias, 1959; Steinman, 1965; Putnam et al., 2005). This subset of cones at the PRL may have functional preeminence, as fixational eye movements re-center the object of interest on this location (Pritchard, 1961). However, previous studies showed that the PRL does not colocalize with structural features such as peak cone density (Putnam et al., 2005) or the center of the foveal pit volume (Wilk et al., 2017a).

With recent optical tools, the mosaic of even the smallest cone photoreceptors in the foveola can now be resolved in the living human eye and visual function such as visual sensitivity can be simultaneously probed psychophysically with cellular precision (Harmening et al., 2014). In a first attempt to better understand how the specific mosaic of cone photoreceptors within the foveal center has functional consequences to vision, we here use this experimental access to study the direct relationship between the cellular makeup of the foveal center and sensitivity to light in four human participants.

## Methods

In four human participants (1 female, 3 males; 35 ± 6 years), the cellular topography of the fovea, light sensitivity to cone-sized stimuli, and the preferred retinal locus of fixation where mapped with cellular precision using an adaptive optics scanning laser ophthalmoscope (AOSLO) microstimulator. The participants were three of the authors (P1= ND, P2 = JLR, P3 = WMH), and one lab member (P4). Pupil dilation and cycloplegia were induced by instilling one drop of 1% Tropicamide 15 minutes before the beginning of an experimental session. For each participant, a custom dental impression (bite bar) was used to immobilize and control the position of the head during imaging and stimulation. Written informed consent was obtained from each participant and all experimental procedures adhered to the tenets of the Declaration of Helsinki, in accordance with the guidelines of the independent ethics committee of the medical faculty at the Rheinische Friedrich-Wilhelms-Universität of Bonn, Germany.

### Adaptive optics scanning laser ophthalmoscope microstimulator

The central ∼1 degree of the right eye of each participant was imaged and targeted test sites were simultaneously stimulated with a custom multiwavelength AOSLO (for details of the system see (Poonja et al., 2005; Grieve et al., 2006; Domdei et al., 2018, 2019)). In brief, the output of a supercontinuum light source (SuperK Extreme EXR-15, NKT Photonics, Birkerød, Denmark) was split into two light channels by serial dichroic and bandpass filtering: one near infrared (IR) light channel was used for imaging and wavefront sensing (840 ± 12 nm, FF01- 840/12-25, Semrock, Rochester, USA), and a visible green light channel for microstimulation (543 ± 22 nm, FF01-543/22-25, Semrock). Adaptive optics correction, run in closed loop operation at about 25 Hz, consisted of a Shack-Hartmann wavefront sensor (SHSCam AR-S- 150-GE, Optocraft GmbH, Erlangen, Germany) and a magnetic 97-actuator deformable mirror (DM97-08, ALPAO, Montbonnot-Saint-Martin, France) placed at a pupil conjugate. Imaging and stimulation beams, traversing the system coaxilly, were point-scanned across the retina, spanning a square field of 0.85 x 0.85 degrees of visual angle. The light reflected from the retina was detected in photomultiplier tubes (PMT, Photosensor module H7422-40 and -50 Hamamatsu Photonics, Hamamatsu, Japan) individually for each channel, placed behind a confocal pinhole (pinhole diameter = 20 µm, equaling 0.47 (840nm) and 0.71 (543nm) Airy disk diameters). The PMT signals were continuously sampled by an FPGA board (ML506, Xilinx, San Jose, USA), producing video frames with 512 x 512 pixel (spatial resolution = 0.1 arcmin of visual angle per pixel) at 30 Hz by combining the PMT’s voltage output with the positional data of the scanners. To increase dynamic range and contrast, the 543 nm stimulus light was modulated by two cascaded acousto-optic modulators (Domdei et al., 2018). The stimulation light was attenuated by 3 log units using a neutral density filter before coupling light into the AOMs, ensuring an optimal operating range of the AOMs. The combined effects of chromatic aberrations of the human eye were minimized as follows: longitudinal chromatic aberration, LCA, was compensated by a static relative vergence difference of 1 diopter between the 840 nm and 543 nm light channel (Atchison and Smith, 2005), and an individual adjustment of defocus of the DM for each eye, prioritizing image quality in the visible channel. Transverse chromatic offsets were compensated dynamically with every stimulus presentation by lateral stimulus location offsets driven by correction signals from Purkinje-based pupil monitoring (Harmening et al., 2012; Domdei et al., 2019).

### Cone density maps and cone density centroid

Continuous maps of cone photoreceptor density were generated by recording a 10 second AOSLO video with an imaging wavelength of 543 nm (P1 and P2) or 788 nm (P3 and P4) while the participant was asked to fixate on a small flashing target in the center of the imaging raster. High signal-to-noise ratio images were then generated by strip-wise image registration and averaging of individual frames of those videos, while manually excluding single frames with failed registration. In these normalized images, one human grader marked the location of each cone, assisted by convolutional neural network custom software (Cunefare et al., 2017; Reiniger et al., 2019). Matlab’s Voronoi function (*voronoiDiagram*, Mathworks, Inc., Natick, MA, USA) was used to compute each cone’s area. Cone density at each pixel of this image was then calculated by first identifying the nearest 150 cones around each pixel in the image. Cone density was then defined by dividing the number of cones by their summed area. The cone density centroid (CDC) was the retinal location found as the weighted centroid (Matlab function *regionprops(Image,’weightedcentroid’*)) of the area containing the highest 20 % cone density values.

### Determination of cone outer segment length

Spectral-domain optical coherence tomography (OCT) images were acquired for all 4 participants in a 5 x 15 degree field, centered on the fovea with a B-scan spacing of 11 µm (HRA-OCT Spectralis, Heidelberg Engineering, Heidelberg, Germany). For further processing, the central 45 B-scans around the foveal pit were selected. Cone outer segment (OS) length was defined as the space between visible bands 2 and 3 in each B-scan, thus is the space between the ellipsoid zone and the interdigitation zone, respectively (Spaide and Curcio, 2011). In a first step, these two bands were segmented by manually adjusting a brightness-based automated detection algorithm (Fig. 5A). Because the retinal pigment epithelium is thought to have a relatively uniform thickness and flat layout, a two-dimensional area fit across all B-scans for the third band was computed, reducing artifacts of the individual band marking. The width of both bands was modelled with a 1D gaussian profile, centered on the band marking. The OS length was defined as the linear space between the steepest parts of the second band’s declining and the third band’s rising slope (see also Spaide and Curcio (2011)). In the final step, the two-dimensional OS length map was smoothed along the orthogonal B-scan axis with a Savitzky-Golay filter to remove residual artifacts of individual band markings between the single B-scans and then rescaled to obtain equal increment per pixel values in all directions. These 2D maps of OS length were registered with retinal AOSLO imagery by centering their maximum value at the CDC.

### PRL determination

The preferred retinal locus of fixation (PRL) was determined by recording the exact retinal location at which a small, flashing visual stimulus landed during attempted fixation. The fixation stimulus was created by modulating the imaging channel of the AOSLO to turn off in a central region of the raster, creating a small visible black square, 1.6 × 1.6 arcmin nominal size, shown against the visible 840 nm, 51 × 51 arcmin scanning background. The stimulus flashed continuously at 3 Hz and its retinal landing positions were recorded in three consecutive 10 second AOSLO videos. Retinal fixation locations were then found by, (1) automatically registering each frame strip-by-strip to a common reference frame of a single video (Arathorn et al., 2007), (2) manual deletion of incorrectly registered frames (usually due to microsaccades, insufficient image quality or eye blinks), (3) tabulating all stimulus locations within the remaining, co-registered frames (Fig. 1A). The final PRL estimate for each eye was the median stimulus location across all three videos, equaling on average 387 frames (Fig 1B). This location was defined as 0 degree eccentricity.

**Figure 1:**
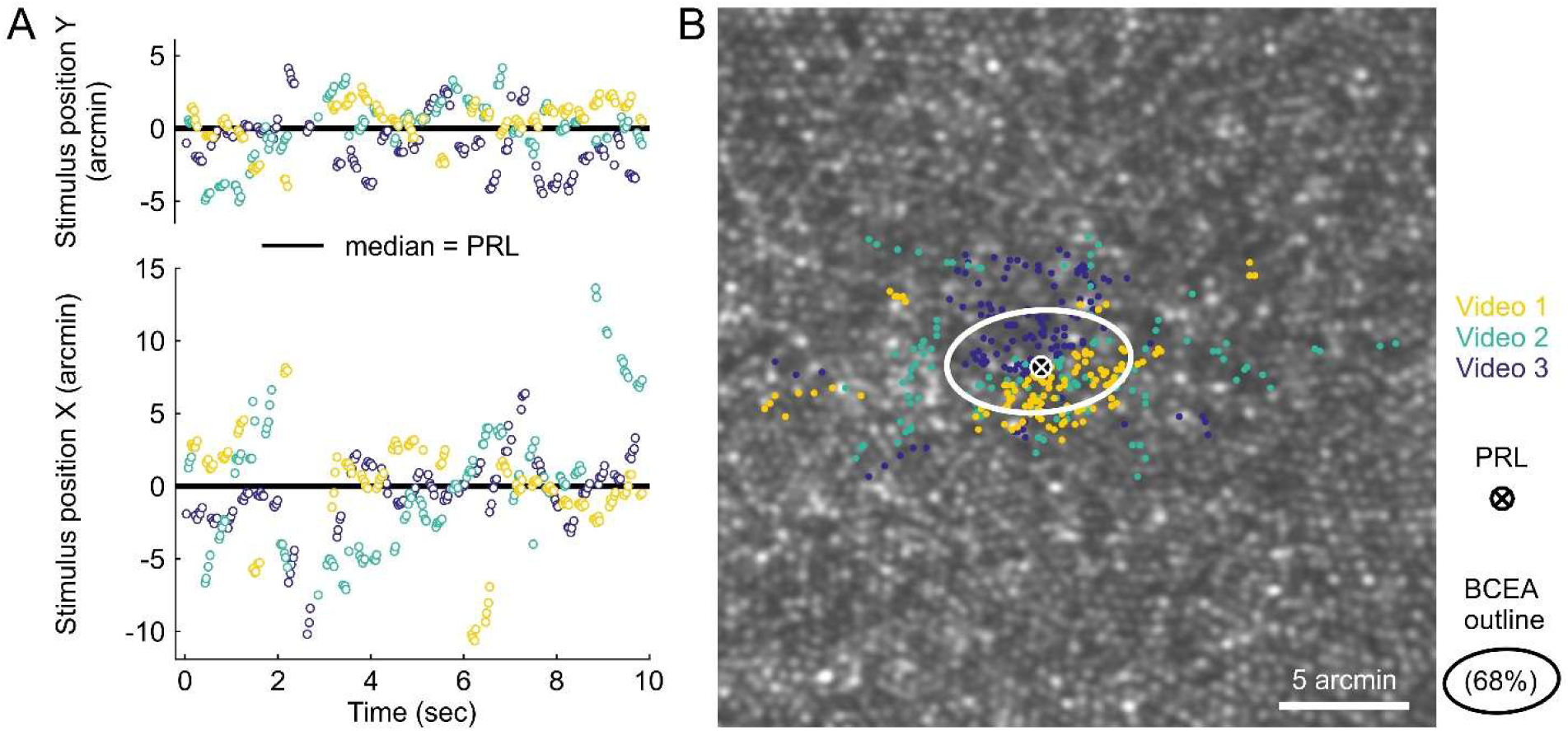
Preferred retinal locus (PRL) determination from three 10 sec videos. A) The nominal fixation target was a 1.6 × 1.6 arcmin square presented at a fixed position in the center of the AOSLO raster flashing with 3 Hz. High resolution eye motion traces were recorded in three 10 second epochs, tracking the position of the target’s center in retinal coordinates. Single dots represent frame-by-frame derived retinal coordinates of the target, colors indicate repeats. B) Stimulus positions in relationship to the foveal cone photoreceptor mosaic. The bivariate contour ellipse was set to contain 68 % of all stimulus locations. The participant’s PRL (white ellipse and marker) was computed as the median data point, pooling all locations of the three consecutively recorded videos.

### Increment sensitivity thresholds

After PRL determination, participants underwent small spot sensitivity testing at multiple sites within their foveola, with the PRL as a spatial anchor at the center of the test locations. Sixteen additional test locations were selected manually around the PRL, close to the intersections of two concentric perimeters (spaced 6 and 12 arcmin) around the PRL with the horizontal, vertical, and diagonal meridians (Fig. 2A and B). At each of the 17 test locations, sensitivity thresholds were determined as the median of 3-5 repeat measurements per location. In each experimental run, three out of the 17 locations that were oriented closely along a vertical axis were tested as a group, with the individual locations tested pseudo-randomly interleaved within the group. This approach minimized participant fatigue, allowed for more regular breaks between runs, and also minimized background intensity changes that are present towards the horizontal edges of the imaging raster (± 0.05 contrast change), because stimuli could be presented in the central part of the imaging raster at all times. Sensitivity thresholds at each test location were estimated with QUEST, an adaptive staircase method (Watson and Pelli, 1983), adjusting stimulus intensity following a Yes/No-test paradigm with logarithmic step sizes (King-Smith et al., 1994). Because of the visible IR imaging raster, stimuli were presented against a reddish background (about 5 cd/m² photopic luminance), rendering our sensitivity testing paradigm to yield increment sensitivity thresholds (see also Harmening et al. (2014); Tuten et al. (2017)). The test stimulus was a green square with 7 pixels edge length in scanning raster coordinates. Considering a residual defocus of 0.03 D (Meadway and Sincich, 2018), the stimulus edge length given by the full width at half maximum was about 0.8 arcmin, or between 150 % (P1) and 180 % (P4) the diameter of the smallest cones. Due to the scanning nature of the AOSLO, where visual stimuli are rendered pixel-by-pixel while the illumination beam traverses the retina, the presentation time spanned 350 µsec from the first to the last pixel of the stimulus. This equals a net illumination time per stimulus of 2.5 µsec. Stimulus presentation progression was self-paced and successful deliveries were accompanied by an auditory cue. Stimulus delivery was blocked if either the online stabilization algorithm or the eye tracking for TCO compensation failed, and the participant had to repeat the trial (∼32.4 % of all 5674 stimulus presentations). Threshold estimation was completed after 12 trials if the standard deviation (STD) value of QUEST was less or equal to 0.10 log arbitrary AOM voltage drive units (Fig. 2D). If STD was higher, the run was extended by additional trials until the STD criterion was met. The run was terminated and had to be repeated if the STD criterion was not met after 18 trials. Runs were repeated until a minimum of three valid threshold estimates per test location were recorded.

**Figure 2:**
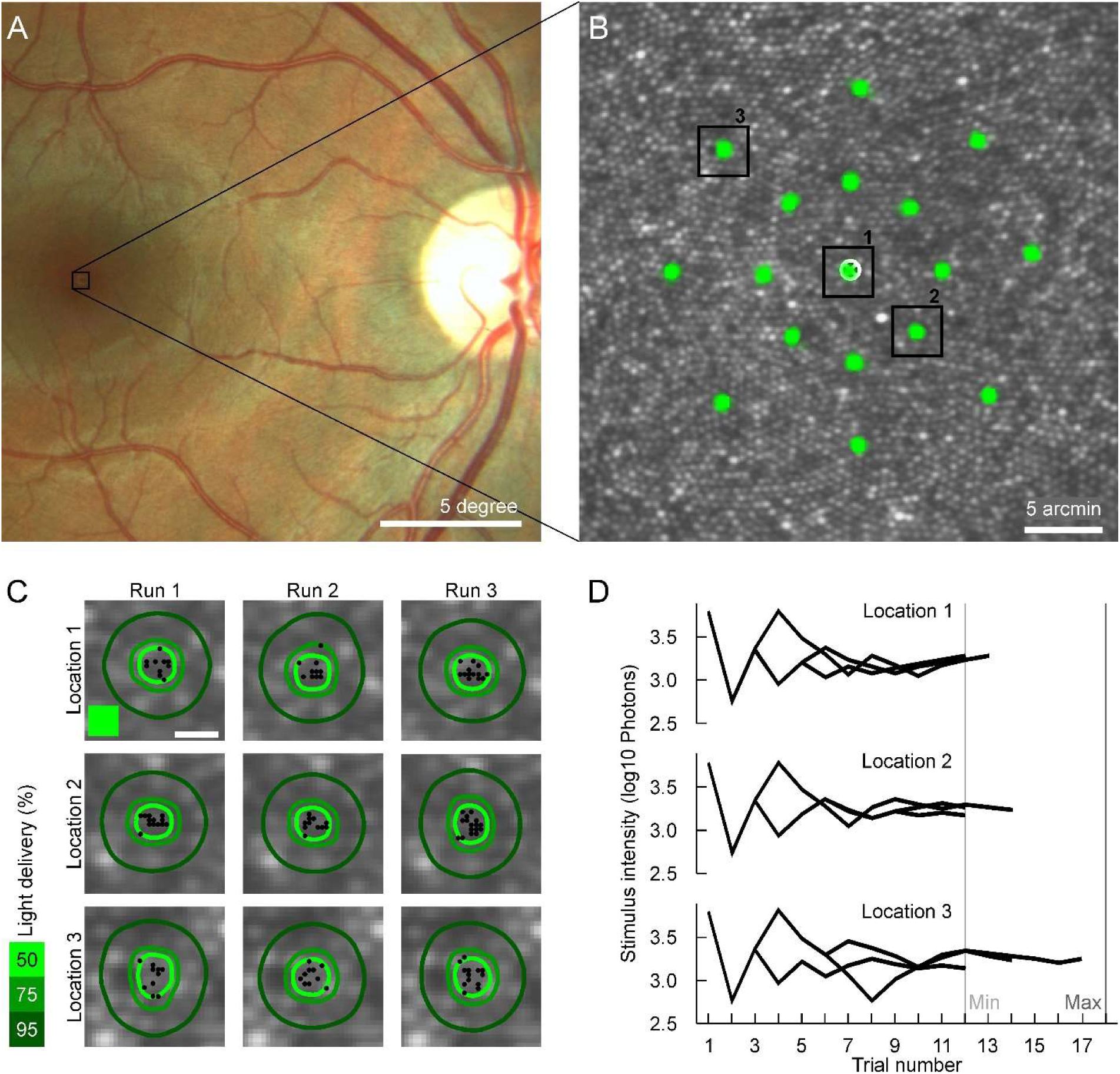
AOSLO-based microstimulation. A) Color fundus image of one participant. The grey square shows the size and position of the AOSLO imaging and stimulation raster on the retina, positioned at the center of the fovea. B) Cropped view of an averaged frame showing the AOSLO image of the central fovea. Retinal stimulation sites are marked by transparent green markers, three sites are highlighted by black boxes to be further analyzed in the next panels. C) Zoomed-in view of selected target sites. Markers indicate individual stimulus locations during repeated stimulation for threshold estimation. The green square represents the stimulus in raster pixel size. The contour lines mark 50 %, 75 %, and 95 % of the summed light delivery for each run. The scale bar is 1 arcmin. D) Exemplary progression of stimulus intensity based on the current threshold estimation via QUEST with three runs per test site. The individual run was completed, when the standard deviation of the estimated threshold was less or equal 0.10, resulting in a varying number of trials per test site and run.

Real-time image stabilization enabled retinal tracking and repeated stimulation of the targeted test sites (Arathorn et al., 2007). In a subsequent offline analysis, trials with suboptimal delivery (more than 0.6 arcmin deviance from median delivery location, equaling about 1.2 cone diameters) were excluded from further analysis. The non-linear swing speed of the resonant scanner produced a non-uniform IR background across the field, with increasing brightness close to the vertical edges of the field. For compensation, the individual trial intensity was corrected based on the position in the imaging raster and the resulting stimulus contrast, with compensation factors ranging between 0.95 and 1.00. After trial rejection and intensity correction, QUEST was computationally re-run with the updated stimulus intensities to recompute the final threshold estimate. Trial deletions on account of stabilization errors produced higher QUEST STDs. In a second step, thresholds with an STD higher than 0.15 log a.u. were excluded from the following analysis. Sensitivity thresholds are finally computed and reported as the number of photons incident at the cornea at the threshold intensity (see next chapter).

### Conversion of arbitrary power units to number of photons at the cornea

Before and after each experimental run, maximum output power of the AOSLO stimulation light channel was measured at maximum AOM drive voltage with a silicon photodiode and power meter (S130C and PM320E, Thorlabs GmbH, Bergkirchen, Germany) in the transmitted portion of the stimulation beam after a 90/10 (Transmisson/Reflection) beam splitter placed in the light delivery arm of the AOSLO. The average of these two measurements was used for actual stimulus power calculation of a given run. Typically, laser power fluctuated by less than 1 % between measurements. With this, the QUEST threshold estimate (ThreshEst.), so far given in log10 arbitrary AOM drive units, could be converted into a number of photons at the cornea using formula (1):

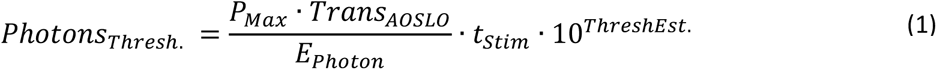

with P(Max) being the maximum AOM output power measured, Trans(AOSLO) being the relationship of maximum power fed into the AOSLO and the power detected at the eye’s pupil position (determined earlier to be 0.065) and t(Stim) being the stimulus duration during a single presentation (2.45 µsec). E(Photon), the photon’s energy, was calculated as:

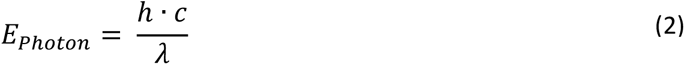

with h, Planck’s constant, c, the speed of light, and λ, stimulus light wavelength (543 nm).

### Modelling of cone light capture and ISETBio

To assess the impact of the exact stimulus location relative to the individual cone mosaic, a custom spatial model of quantal catch in the targeted cones was implemented in Matlab, similar to (Harmening et al., 2014). First, all stimulation trials were registered to a high-signal-to-noise image of the foveal center used to create the cone density map. Second, cone center locations were used to compute a complete Voronoi tessellation of the mosaic defining the inner segment area of the individual cones. Third, the actual absorption characteristic across this inner segment area was modelled by a two dimensional gaussian with a sigma value creating an aperture of 0.48 of the equivalent diameter (Macleod et al., 1992). The retinal stimulus was a two-dimensional convolution of the initial 7-by-7 AOSLO raster pixel stimulus and a diffraction limited point spread function with a residual defocus of 0.03 D (Meadway and Sincich, 2018). In the final step, the convolved stimulus was multiplied by the cone absorption matrix to arrive at the amount of stimulus light that was absorbed by each cone. The reported value of percent of light per cone is based on the total light distribution given by the convolved stimulus matrix. But this model lacked a more generalized testing of the impact on thresholds by variations in the OS length or cone density. To test the hypothesis that the cone’s biophysical properties such as cone density, cone diameter and cone outer segment length had an impact on sensitivity thresholds, the Image Systems Engineering Toolbox for Biology (ISETBio) for Matlab (Cottaris et al., 2019, 2020), was used. This computational-observer model simulates the cone photoreceptor isomerizations and photo currents based on physiological constraints. To set up ISETBio, the following parameters were used: the ‘scene’ was a 511 x 511 pixel image with the central 7 x 7 pixel containing the stimulus. The field of view was set to 0.85 degree and the background luminance set to 0.1 cd/m^2^. The wavefront was set to be diffraction limited, with a varying residual defocus of 0.01, 0.03, and 0.05 diopters and zero LCA. The hexagonal mosaic function was used with a custom cone spacing and optical density. The optical density was calculated as the product of the OS length and an average absorbance of 0.014 µm^-1^(L-cones = 0.013 ± 0.002 µm^-1^and M-cones = 0.015 ± 0.004 µm, (Bowmaker et al., 1978)). Each condition (varying cone density, optical density and both coupled) was repeated 10 times taking simulated neural noise of the photoreceptor cells into account. To model different stimulus positions relative to the mosaic, the stimulus was shifted in steps of 0.6 arcsec (pixelwise at a magnification of 10). The test site’s cone class composition was controlled by adjusting the L- and M-cone spatial density parameter for each cone class and carefully checking the center surround configuration of the generated mosaic. These two simulations were both run with a residual defocus of 0.03 and repeated 33 times per condition. For Figure 10, the average of these data sets was plotted with the error bars reflecting one standard deviation. The slopes found by the ISETBio model for each parameter were used to calculate a factor specific offset ΔT_i_ from the average threshold for each of the test sites:

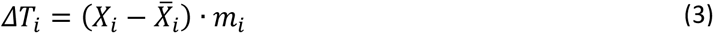

With X_i_ being the test site’s value of the factor *i* (e.g. OS length), 𝑋 the factor’s average (31.39 µm) and m_i_ the factor specific slope. The sum of all three ΔX_i_ values yielded the test site’s threshold offset from the average threshold. This way it was possible to compare the observed thresholds with an expected threshold based on the models without knowing the intercept of the factor specific function.

## Results

### Retinal topography and the PRL

With AOSLO imaging, the foveal mosaics of all four observers could be resolved, and all cone photoreceptors could be identified and marked to create continuous topographical maps of cone density. Participant numbering, P1-P4, was ordered for all following analysis according to their peak cone density values (13733, 15230, 18023, and 18406 cones/degree² for P1, P2, P3, and P4 respectively). At the location of the cone density centroid (CDC, see Methods) the density values were slightly lower P1: 13460, P2: 15199, P3: 16956, P4: 16914 cones/deg². Cone density dropped rapidly with increasing eccentricity. Across subjects, cone density dropped on average to 66 % of the peak cone density at a distance of about 20 arcmin from the CDC. However, there were marked differences in topographical profiles between subjects. For instance, P1 had a rather plateau like cone density distribution, while P4 showed a steep decline with a two-pronged cone density profile. The other participants had a similar slope in cone density profiles.

The two-dimensional map of cone outer segment (OS) length as determined by high-resolution OCT imaging was found to show a similar topography as cone density in each eye. P1, the participant with the lowest peak cone density, also had the lowest maximum OS length of 28 µm. For P2, P3, and P4 the maximum OS length was 28, 32, and 40 µm, respectively. The range of OS length was 3 µm (25 to 28 µm) for P1, 4 µm (28 to 32 µm) for P2, 4 µm (29 to 33 µm) for P3, and 7 µm (32 to 40 µm) for P4 within the central 12 arcmin radius around the CDC. The PRL was found to be offset from the CDC and the location of highest OS length in all eyes. It was shifted away from a nasal-inferior quadrant of the retina with respect to the CDC with an average offset of about 3 arcmin. For each participant, the offset, as well as the angle, were different, ranging from 1 arcmin and 15 degree (P1) to 3 arcmin and 178 degree (P4) with a good repeatability of about 1 arcmin (Fig. 3A). PRL ellipses were found to be small in all participants (BCEA: P1 = 50; P2 = 67; P3 = 16; P4 = 30 arcmin²), indicative of high fixation stability, a prerequisite for the following cone-targeted sensitivity testing.

**Figure 3:**
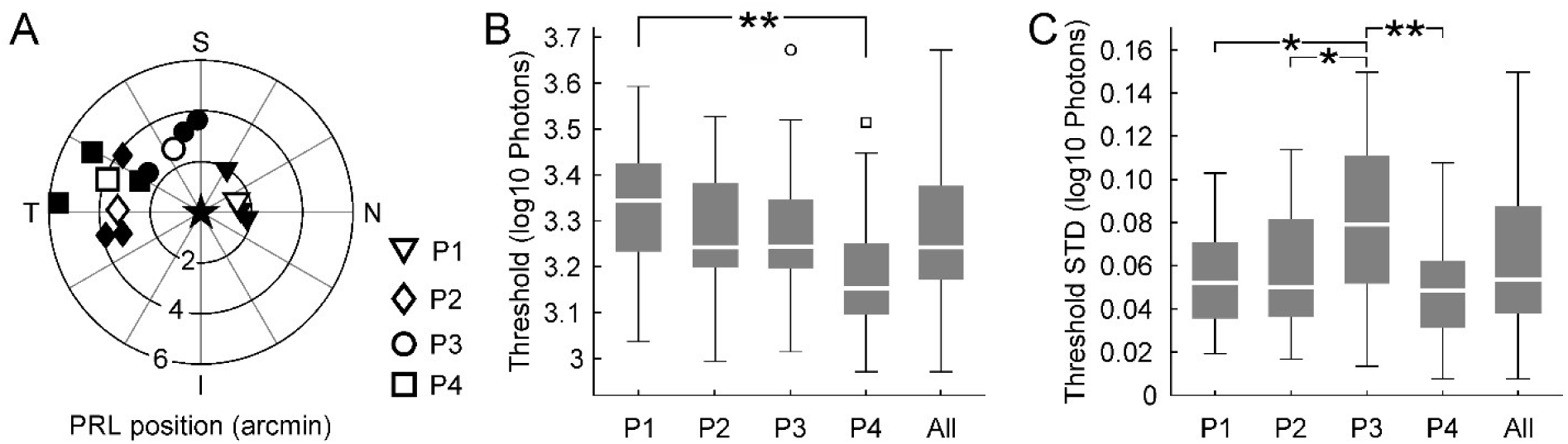
Fixation behavior and foveal increment sensitivity. A) The PRL of each participant (P1-4) is plotted relative to the location of the cone density centroid (plot origin, ‘star’ marker). Filled markers represent single measurements, open markers are the final PRL based on three repeat measurements. Letters marking Nasal, Temporal, Inferior, and Superior. B) Pooled sensitivity thresholds across all test sites for each participant separately (P1-P4), and combined (“All”). While the range (∼0.5 log10 units, whiskers), as well as the highest and lowest thresholds, were very similar across participants, the median, 1st, and 3rd quartile differed. C) The observed standard deviation (STD) of repeated threshold estimations at the same test site was similar for three participants. Variability for P3 was higher than for the other three. 85 % of all observed STDs were less or equal to 0.10 log10 photons.

### Small spot sensitivity

Small spot sensitivity thresholds across subjects were found to lie between 2.97 and 3.67 log10 photons, spanning an overall range of 0.7 log10 photons. The combined median of all thresholds across retinal locations and participants was 3.25 log10 photons at the cornea and the average 3.27 ± 0.16 log10 photons. While the range of thresholds across the four participants was similar (3.04 to 3.59, 2.99 to 3.53, 3.02 to 3.67, and 2.97 to 3.51 log10 photons for P1, P2, P3, and P4 respectively; see Figure 3B), there was a continuous shift of the median, 1st, and 3rd quartile of thresholds from P1 to P4. The median thresholds shifted by 0.2 log10 photons from 3.35 for P1 towards 3.15 for P4 (3.24 for P2 and P3). For P2 and P3 the 1st and 3rd quartile of thresholds were lower than for P1, but higher than for P4, this difference was not significant (Q1: 3.23, 3.20, 3.20, and 3.10; Q3: 3.43, 3.38, 3.34, and 3.25). The repeatability for each test site, computed across three to five reruns, was high, with an average standard deviation of 0.06 ± 0.03 log10 photons (Fig. 3C). The standard deviation across reruns was higher for P3 (average = 0.08 log10 photons) compared to the other three participants (0.05 log10 photons).

### Correlation between retinal structure and function

To bring sensitivity thresholds into spatial correspondence with the structural data and fixation behavior, averaged retinal images derived from all four independent analysis steps (cone density maps, OS length map, PRL determination, and microstimulation target sites) were carefully aligned with each other (Fig. 4 and 5), to allow a pointwise and cellular resolved comparison between retinal structure and function.

**Figure 4:**
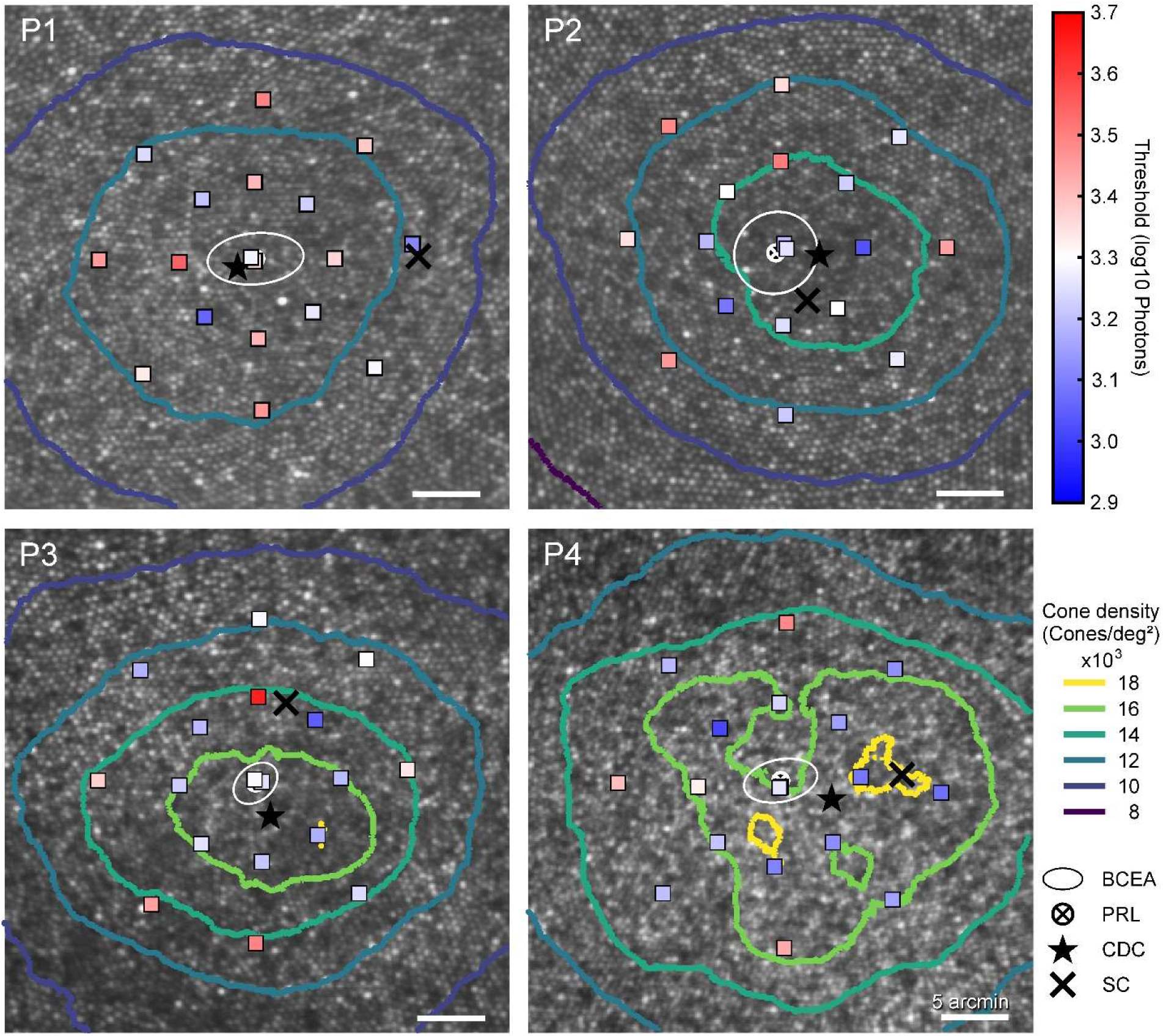
Sensitivity in relationship to cone topography and fixation behavior in all participants. Squares mark the average stimulus location for each test site, coloring reflects the median sensitivity threshold in log10 photons. Marker area is four times the actual stimulus area on the retina and equals the average 95 % outline of the summed light delivery (see Fig.1C). Cone density is indicated by colored contour lines. The PRL (white circle and crosshair), the location of the cone density centroid (star) and an estimated location of the sensitivity centroid (SC, cross) are shown as well.

**Figure 5:**
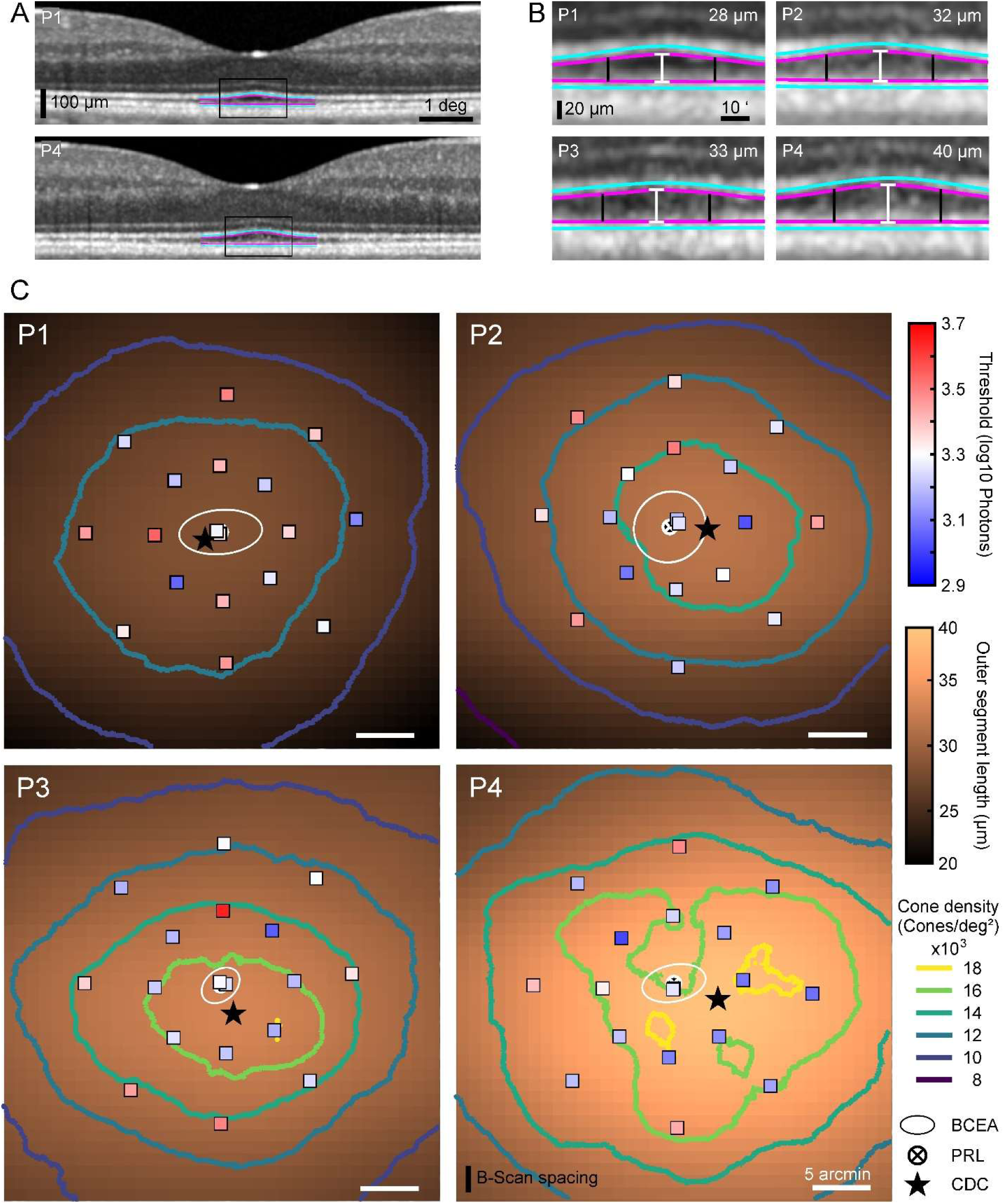
Cone outer segment length topography. A) Foveal OCT B-scans from P1 and P4, and a zoomed-in view of the OS in the central fovea. B) Further zoomed-in view of central OS and measurement demonstration for all 4 participants. The band’s center (cyan line) was semi-automatically marked. The OS start and end (magenta) was found by fitting a 1D gaussian centered on the 2nd and 3rd band. The estimated maximum OS length is stated in the upper right corner. C) Interpolated 2D map of cone OS length in the central fovea for each participant. The black vertical lines in (B) indicate the region of the 2D presentation in (C).

As a first observation, for none of the participants, the target site with the lowest threshold and therefore highest sensitivity was at the PRL or fell within the fixation ellipse (5.7, 6.5, 6.5, and 6.0 arcmin distance). The average distance between the test site with lowest threshold and highest cone density was 7.3 arcmin. In P1 and P2 the CDC was closer to the target site with the lowest threshold than the PRL (CDC distances: 4.5, 3.3, 7.8, 9.8 arcmin).

The range of cone densities at the test sites differed clearly within and between participants, with a difference from minimum to maximum of 2300 cones/deg² (11215 to 13198 cones/deg²) for P1, 3400 cones/deg² (12564 to 14971 cones/deg²) for P2, 6100 cones/deg² (11786 to 17881 cones/deg²) for P3, and 3800 cones/deg² (14364 to 18132 cones/deg²) for P4. While the highest cone density of P4 was about one third higher than the peak cone density of P1, sensitivity thresholds were similar at those locations. However, a general trend was noticeable that most of the sensitivity thresholds observed in P1 were higher than most of the thresholds of P4.

For further correlation analysis, we used the median threshold of repeated threshold estimations at the same test location. The difference between individual sensitivity thresholds and their median can be seen comparing Figure 6A with 6B. While there was no general significant correlation between the distance from PRL and sensitivity thresholds (Fig. 6B), the lowest thresholds were observed at 6 arcmin PRL distance for all participants, and the median threshold at this eccentricity was almost identical (P1 and P2) or even lower (P3 and P4) than the median threshold at the PRL. For the 12 arcmin eccentricity, we observed a similar or higher median threshold compared to the median PRL threshold. In P1 we observed an extremely high threshold at one location that was 0.3 log10 photons higher than the second highest threshold. Plotting the individual thresholds as a function of the test site’s cone density revealed a tendency towards lower thresholds for higher cone densities (Fig. 6C), with a correlation coefficient of ρ = - 0.09 for P1, ρ = -0.49 for P2, ρ = -0.32 for P3 and ρ = -0.52 for P4. The slope showed a tendency to be steeper for higher correlation coefficients: m = -0.02, -0.06, -0.03, and -0.07 log10 photons per 10³ cones/degree² for P1, P2, P3, and P4, respectively).

**Figure 6:**
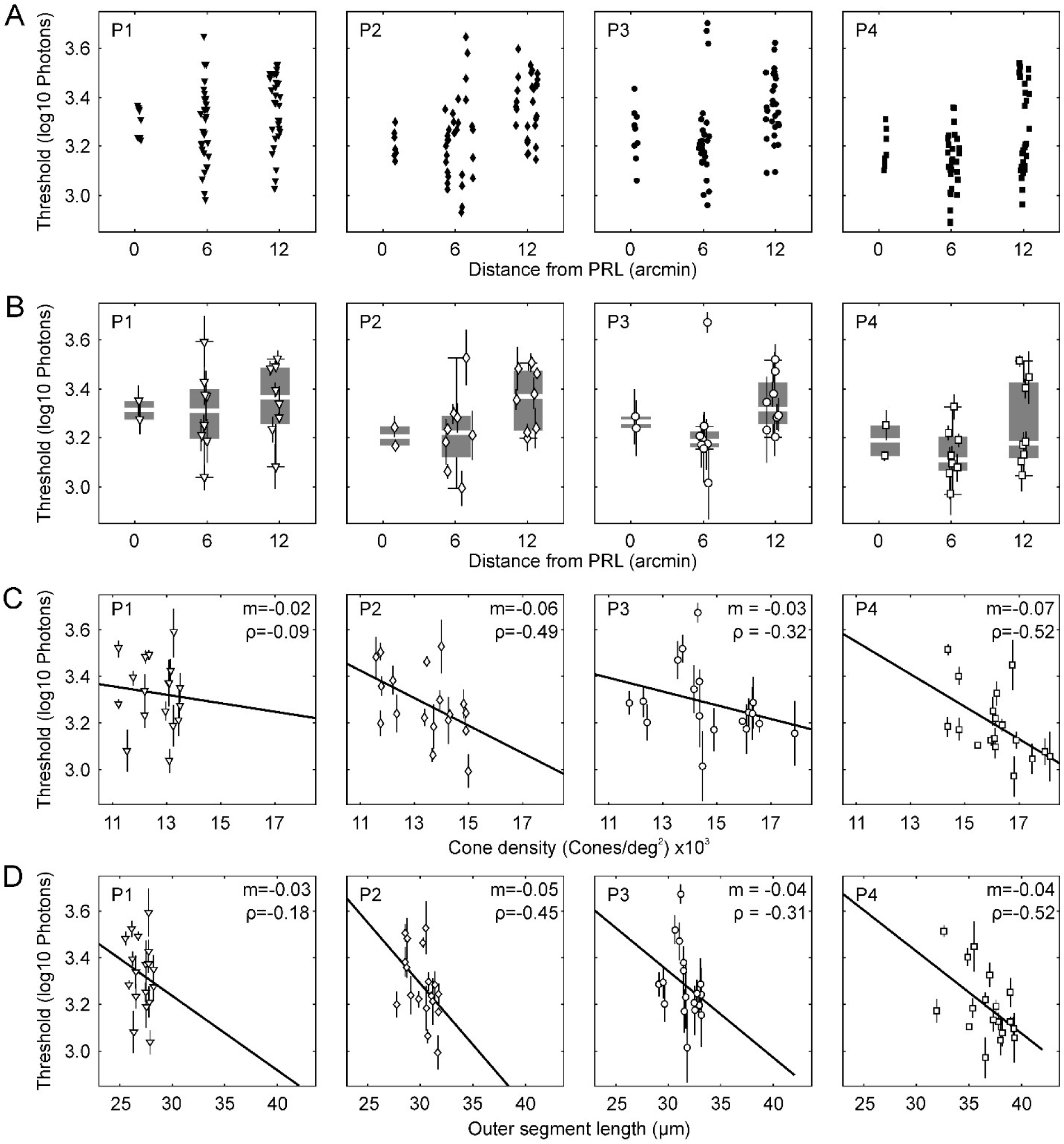
Sensitivity thresholds for each participant in relation to distance from PRL, cone density and outer segment length. A) Individual thresholds for each test site’s distance from the PRL are marked with filled symbols. B) For each test site we used the median threshold (open symbols) of repeated testing (3-5), and reported this value as the test site’s sensitivity. Boxplots show median (white line), first and third quartile (box), whiskers extend to 1.5 fold the distance between the first and third quartile. C) Median thresholds in relation to the test site’s cone density. D) Median thresholds in relation to the outer segment length at the test site as obtained from the 2-D map shown in Fig. 5. Vertical lines in B, C, and D show the standard deviation across repeated runs, thick black lines are linear fits to the data.

Correlating thresholds with foveal cone OS length revealed a similar trend: thresholds were generally lower at sites with higher OS length (Fig. 6D). Correlation coefficients were ρ = -0.18 for P1, ρ = -0.45 for P2, ρ = -0.31 for P3 and ρ = -0.52 for P4. The slope was almost the same across the four participants (m=-0.03, -0.05, -0.04, and -0.04 log10 photons per micron for P1, P2, P3, and P4, respectively).

When data was pooled across subjects, the correlation between the thresholds and the distance from the PRL was given by a slope of 0.65 log10 photons per degree (ρ=0.27, Fig. 7A). The observed decrease of thresholds at 6 arcmin distance from the PRL remained in the combined dataset. Thresholds as a function of cone density had a negative slope of -0.04 log10 photons per 10³ cones/degree² (ρ= -0.45, Fig. 7B). The correlation between thresholds and OS length indicated an average decrease of thresholds by -0.02 log10 photons per micron in the pooled data set (ρ= -0.43, Fig. 7C).

**Figure 7:**
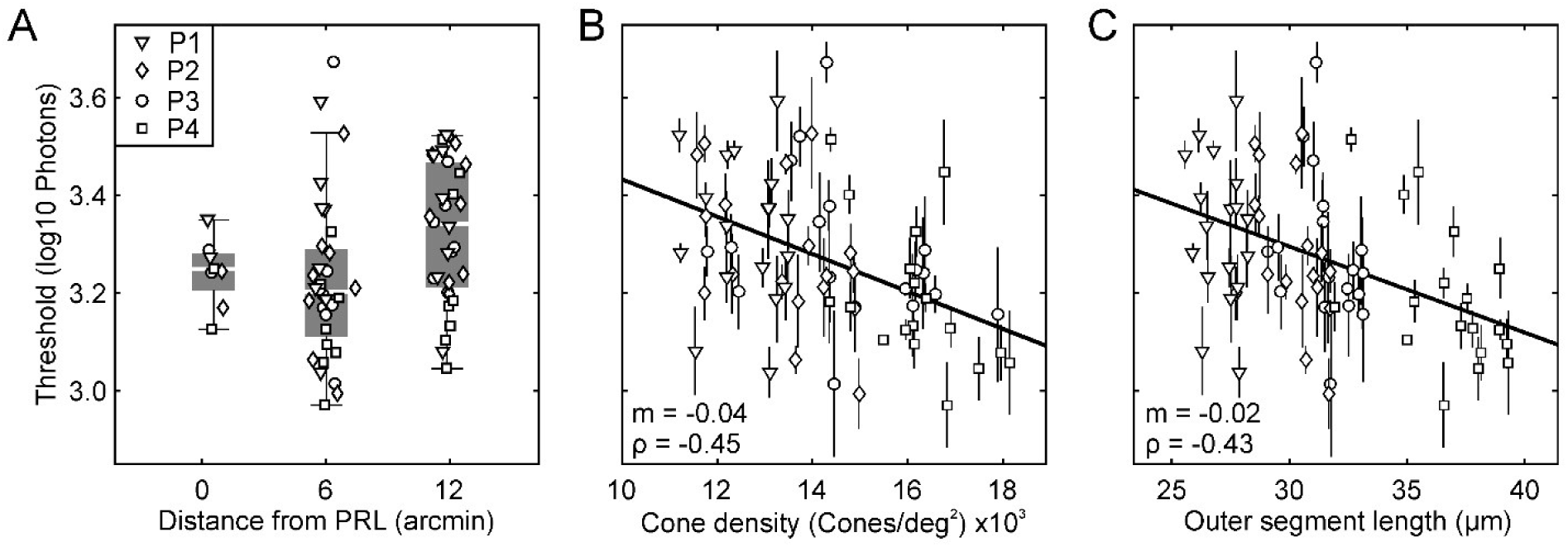
Pooling threshold data across subjects. A) Sensitivity thresholds of all participants as a function of test site distance from PRL, given as median (white line), first and third quartile (box). Whiskers extend to 1.5 fold the distance between the first and third quartile. For the pooled data set, a bend towards lower thresholds at 6 arcmin was observed. B) Thresholds as a function of local cone density at the test site, vertical lines show the standard deviation of threshold estimates for repeated testing. Thresholds and cone densities showed a moderate correlation (ρ = -0.45). C) Thresholds as a function of cone outer segment lengths showing a moderate correlation (ρ = -0.43). All independent variables (Distance, Density, OS length) were pairwise significantly correlated with each other (p < 0.001, data not shown).

Because the individual factors, distance from PRL, cone density and OS length are all highly significantly correlated with each other (p < 0.001, data not shown), a physiological model of cone light capture (ISETBio) was employed to model the impact of two of the three factors independently.

### Modelling the impact of cone density, OS length and distance from PRL on sensitivity

The first hypothesis tested with ISETBio modelling was if the highly significant correlation between detection thresholds and cone density could be caused by spatial summation effects. The stimulus used in sensitivity testing was about 1.5 times the average foveal cone diameter (see methods) and therefore the average distance to the surrounding neighbors could have played an important role. Using the cone spacing according to the individual cone densities of our participants, the model showed a correlation of about -0.01 log10 photons per 10³ cones/degree^2^, roughly four times lower than the observation in the behavioral data (Fig. 8A). ISETBio was also used to test the influence of outer segment length and associated optical density on sensitivity. The outer segment length was estimated for each participant from foveal OCT B-scans (Fig. 5), with a range of 25 µm to 40 µm, and fed into the model. A strong impact of outer segment lengths on thresholds was predicted (ρ = -0.99), with a slope of -0.008 log10 photons per µm OS length (Fig. 8B).

**Figure 8:**
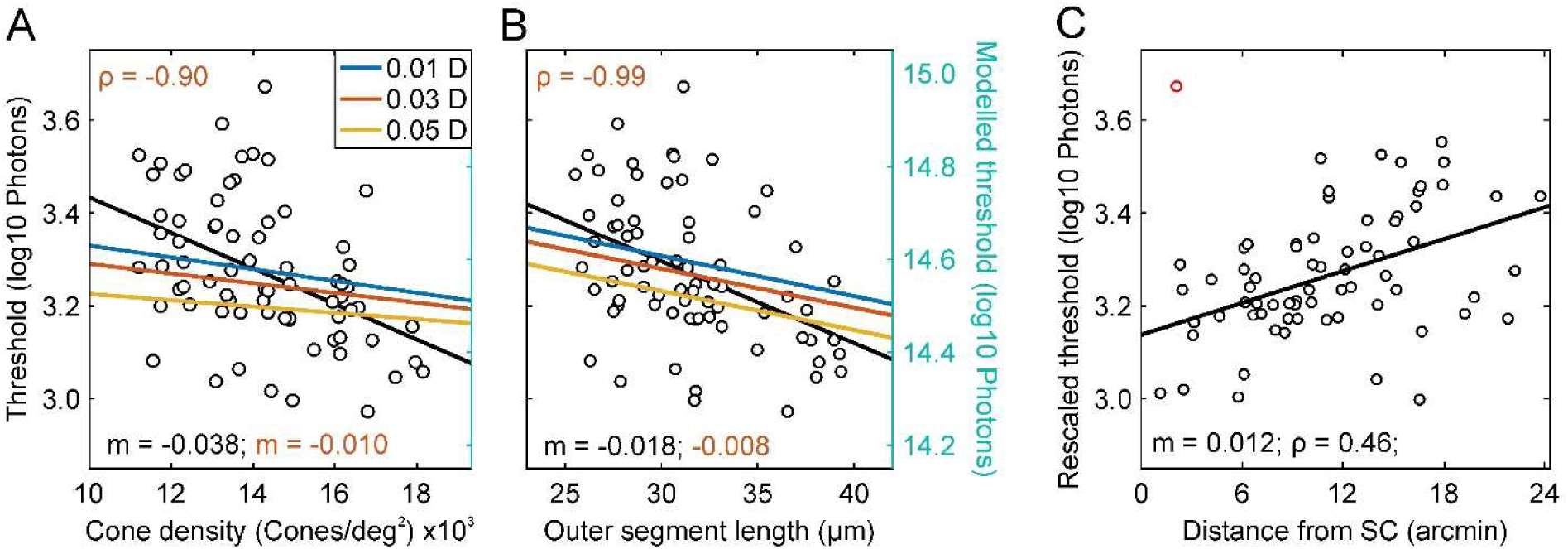
ISETBio model testing the impact of cone density and outer segment (OS) length. A) The ISETBio model evaluating the threshold photons to elicit a certain number of isomerizations, fitted for 0.01 D, 0.03 D, and 0.05 D residual defocus. The simulation revealed only a small influence of cone density and therefore spacing on sensitivity thresholds (m = -0.010 and ρ = -0.90 with 0.03 D), too small to explain the observations. B) To test the influence of outer segment length and therefore optical density on sensitivity, we fed the observed range of outer segment lengths (25 µm to 40 µm) in our ISETBio model, which predicted a strong impact of different outer segment lengths on thresholds (m = -0.008 and ρ = -0.99 with 0.03 D). C) The pooled data of the rescaled thresholds, which had the OS length and density influence removed (see methods), against the distance from the participant’s sensitivity centroid (see Fig. 4). The red marked threshold was ignored in this analysis due to its uncertain nature.

In the following step, the slopes derived for 0.03 D residual defocus (Meadway and Sincich, 2018) were used to remove the estimated proportional influence of cone density and OS length from the observed thresholds. The corrected thresholds were supposed to vary only due to retinal eccentricity. These thresholds were then used to compute the location of the sensitivity centroid (SC) for each participant (Fig. 3) by finding the retinal coordinate yielding the highest value of ρ for corrected thresholds against distance from this coordinate (Fig. 8C). The SC was always offset from the PRL (11.7, 3.8, 4.8, and 7.5 arcmin distance) or CDC (13.1, 2.2, 6.8, and 3.7 arcmin distance). When all rescaled thresholds were plotted as a function of distance to the SC, a linear fit with a slope of 0.69 log10 photons per degree eccentricity or 0.012 log10 photons per arcmin (ρ = 0.46) emerged.

Finally, we applied these three slopes to model the expected shift from the average observed threshold, due to the test site’s cone density, OS length and SC distance (Fig. 9). Because the y-intercept *b* was unknown, the expected shift was calculated for each of the three factors based on the difference from its average. The estimated thresholds were best fitted by a linear regression with a slope of 1.03 (ρ= 0.38, R² = 0.37), confirming that the interaction of cone spacing, OS length and eccentricity yielded a good predictor for sensitivity thresholds. Given a residual variability of ±0.15 log10 photons from the prediction, we also looked at the variability that is possibly due to the exact stimulus position on the cone mosaic. As our stimulus was roughly 1.5 foveal cone diameters in size, the number of isomerizations elicited could have depended on the stimulus being centered on a single cone or in the middle between three cones.

**Figure 9:**
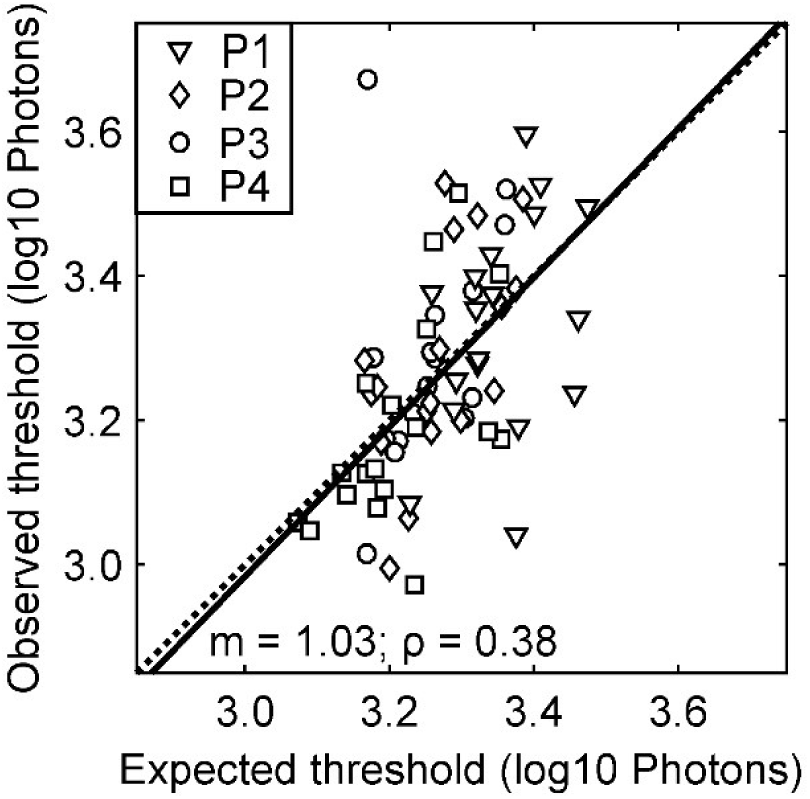
Predicting sensitivity thresholds from retinal factors. The expected threshold was derived from the test site’s cone density, OS length and distance from the sensitivity centroid (Fig. 4). Each parameter was converted into a threshold offset from the average. The solid line is the linear fit to the data, the dotted line shows the 1:1 relationship.

### Modelling the impact of stimulus position and cone class composition

To test the hypothesis that the exact placement of the test stimulus relative to the cone mosaic bears on sensitivity at that site, photon catch of each cone for the average stimulus location during each threshold experiment was modelled. The exact cone locations relative to the stimulus location were determined by registering the experiment’s video data with the high signal-to-noise image used for cone annotation. An example of the result of that model is shown in Fig. 10A for two different cases. The first case shows two different target sites in the retina of P2. At both target sites, the actual stimulus location on the cone mosaic was similar, but thresholds differed significantly by 0.36 log10 photons (p=0.03, Mann-Whitney U-test, n = 4). In the second example from P4, the stimulus placement differed, the first one was centered on a single cone while the second one was placed in the middle of three cones. For this case we found similar sensitivity thresholds, with a non-significant difference (p=0.49, Mann-Whitney U-test, n = 4). When all thresholds were plotted against light catch in the nearest cone, no significant correlation emerged (Fig. 10B). The ISETBio model, creating a generalized perfect hexagonal retinal mosaic, supported this observation, when shifting the stimulus systematically from a cone-centered position to a position in the middle between cones (Fig. 10C). For such shifts, the resulting change given by the number of isomerizations was 0.1 in log10 space.

**Figure 10:**
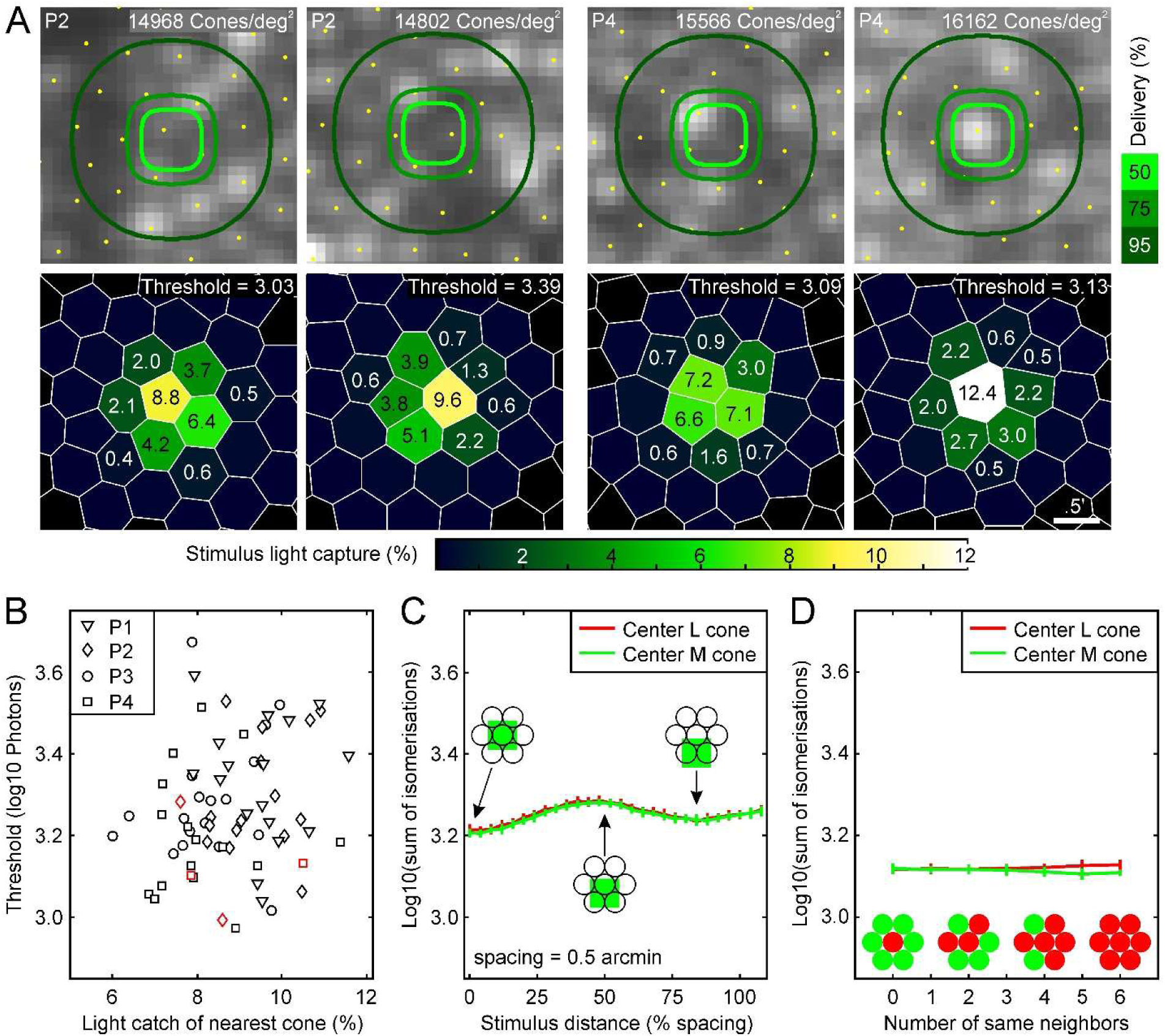
Possible sources of threshold variability. A) We used the cone maps to model the light catch for each cone at the target site. The left two panels show two different test sites of P2, with a similar light catch situation. The observed thresholds for these test sites differed significantly (p=0.03, Mann-Whitney U-test, n = 4). The right two panels show two different test sites of P4, with different light catch situations. In the first condition three cones are supposed to catch the same number of photons, while in the second condition the major portion of stimulus photons are being caught by a single cone. The observed thresholds for these test sites differ insignificantly (p=0.49, Mann-Whitney U-test, n = 4). B) Pooling the data across all participants. Small light catch numbers indicate a 3-cone position, high numbers a single cone center position. Example data sets shown in A were highlighted with a red marker outline. There was no significant correlation between observed thresholds and stimulus delivery condition. C) This observation was confirmed by an ISETBio model testing the influence of stimulus position on the number of isomerizations. The model showed a maximum change of 0.1 log10 isomerizations for different stimulus positions. D) ISETBio model to test the influence of different cone class compositions at the test site. This model does not contain any inter cone class inhibition and therefore shows only a slight increase of isomerizations if solely L-cones were activated. The error bars in C and D indicate the ISETBio simulated retinal noise.

There was no difference if the center cone was a L- or M-cone. Such small changes could not explain the observed variability of ±0.3 log10 photons, but could be one reason for the observed intra-run variability of 0.06 log10 photons due to small stimulus displacements caused by residual errors of the real-time stabilization (Fig. 1C). Another reason causing such variability could be the composition of cone classes at the stimulus location. Again, ISETBio modelling showed only minor changes in the sum of isomerisation for different cone mosaic compositions (Fig. 10D). The maximum difference was 0.02 log10 isomerisations for a stimulus location consisting of only L-cones versus only M-cones. As the actual composition of the cone mosaic in our four participants was unknown, this model did not contain any specific L-M-cone interactions and only used the cone class specific spectral sensitivity.

## Discussion

The human fovea is the result of morphologic specializations culminating in tight photoreceptor packing and elongation of the photopigment containing outer segments (OS). Eye movements constantly align fixated objects of interest to a distinct group of cones in the fovea, the preferred retinal locus of fixation (PRL). The PRL and the retinal location where anatomical features peak are known to be offset from each other. By using an adaptive optics scanning laser ophthalmoscope (AOSLO) as high-resolution imaging and microstimulation platform, the functional profile of the foveola was further investigated. In four observers, increment sensitivity thresholds at a number of target sites around the PRL were correlated with retinal factors such as cone density, photoreceptor OS length, and the distance from the PRL. We generally found a plateau-like sensitivity profile within the central 6-arcmin radius around the PRL with a declined sensitivity at about 12 arcmin eccentricity, resulting in a much flatter slope than the anatomical topography of cone density and OS length changes in the same region. However, about 37 % of the variability in thresholds could be explained by a combined model of the impact of cone density, OS length and distance from the PRL on thresholds, indicating that the exact makeup of the foveal cone mosaic and fixation behavior are indeed factors of visual function at cellular scale.

The average threshold for detecting a cone-sized visual stimulus against a dim red background (5 cd/m² at 840 nm) was 3.27 ± 0.16 log10 photons at the cornea (Fig. 4). While rods enable the detection of individual photons (Sakitt, 1972; Tinsley et al., 2016), the absolute detection threshold at the fovea, and therefore based on cone vision only, was reported to be about 2.8 log10 photons (Marriott, 1963; Geisler and Davila, 1985) or 2.3 log10 photons (Koenig and Hofer, 2011). Because the here reported thresholds are increments against the IR scanning background, the cones are adapted to the background resulting in elevated thresholds. The best comparison is given by a previous AOSLO microstimulation study on the foveal cone summation area, reporting an average increment threshold of 4.4 log10 photons (Tuten et al., 2018). Because they used a 795 nm wavelength background, their increment thresholds are more elevated. Based on Lamb’s equation (Lamb, 1995) in combination with the cone’s specific spectral absorbance (Schnapf et al., 1987) the sensitivity difference between 840 nm and 795 nm is about 1.1 log10 units, which is very close to the observed difference of the here reported thresholds compared to Tuten et al. (2018).

### The factor “Distance from PRL”

Early studies investigating fixational eye movements reported an “optimal locus”, a small group of cones that is mainly used to resolve any object of interest (Barlow, 1952; Cornsweet, 1956; Steinman, 1965). It was assumed that this retinal location colocalizes with the location of peak cone density. Scanning laser ophthalmoscopy enabled a direct study of the retinal locus used, for example during reading (Mainster et al., 1982). In combination with adaptive optics, it was observed that this optimal locus - now termed “preferred retinal locus”, PRL - was displaced from the location of peak cone density (Putnam et al., 2005). Continuing investigation of the PRL showed that it furthermore did not colocalize with the bottom of the foveal pit or the center of the foveal avascular zone (Bedell, 1980; Zeffren et al., 1990; Wilk et al., 2017b).

In clinical research, the term PRL refers to a newly formed stable location on the retina used for fixation (Crossland, 2011), due to a central scotoma (von Noorden and Mackensen, 1962; Timberlake et al., 1986, 1987; Whittaker et al., 1988) caused by macular diseases (Crossland et al., 2005), such as age-related macular degeneration (Rees et al., 2005). As well as for healthy eyes, the underlying processes of PRL formation are yet unknown and the newly formed PRL is usually a suboptimal retinal location, given the fact that other intact locations of the retina would have provided better acuity (Bernard and Chung, 2018) or higher contrast sensitivity (Rees et al., 2005).

One of the hypotheses we assessed here, was that visual sensitivity contributes to the formation of the PRL. The overall observed fixation stability, given by the BCEA, ranged between 16 arcmin² and 67 arcmin² in our study. This is smaller than previously reported BCEA values between 110 arcmin² and 630 arcmin² measured with SLO (Crossland and Rubin, 2002), presumably due to the increased measurement accuracy with an AOSLO.

We found that, in all eyes, the location of the individual best sensitivity, as well as the empirically determined SC was offset from the PRL (Fig. 3 and 5). However, the distance between the PRL and the SC was small, about 7 arcmin on average, and given the observation that thresholds at the PRL and the 6 arcmin eccentricity were similar, we would agree with the conclusion provided by Putnam et al. 2005, that during natural viewing and the typical blur induced by the eye’s optics, the visual system is insensitive to such subtle offsets, leading to a PRL formation as close as possible to the optimal location.

### The factor “Cone density”

We found a moderate correlation between sensitivity thresholds and cone densities at the tested retinal locations within the fovea. Recent research on light propagation in cone photoreceptors would predict a similar outcome, but more clinically oriented small spot sensitivity testing comes to different conclusions. Modelling light propagation in cone inner and outer segments revealed that a smaller cone aperture is beneficial for increased quantum catch due to the cone’s waveguiding properties (Meadway and Sincich, 2018). Thus, if everything else was held equal, a more densely packed cone mosaic would result in smaller foveal cones and therefore increased sensitivity. In our experiments, the diffraction limited stimulus diameter (0.8 arcmin FWHM) was larger than the individual foveal cone aperture (about 0.5 arcmin), favoring additional summation effects between cones straddling the incident light patch.

In general, spatial summation is closely related to the field size of parasol ganglion cells (Volbrecht et al., 2000). Only within the foveola, spatial summation was found to be smaller than the field size of parasol ganglion cells, but still larger than the field size of midget ganglion cells, with a diameter of Ricco’s area of 2.5 arcmin (Tuten et al., 2018). While the private line forming midget ganglion cells (Polyak, 1941; Boycott and Dowling, 1969) show a relatively stable dendritic field size up to an eccentricity of 4 degrees (Dacey, 1993), the dendritic field size of parasol ganglion cells enlarges rapidly, especially within the 5 degree eccentricity diameter (Dacey and Petersen, 1992). Because the detection of small dim stimuli seemed to rely on parasol ganglion cells, showing a steep increase of summation based on their dendritic field size and the cone’s inner diameter (Curcio et al., 1990; Scoles et al., 2014), and the improved light absorption of smaller cones, a sharp peak of sensitivity within the foveola, centered on the cone density distribution should be expected. Early light sensitivity testing in a perimetry apparatus revealed such a light sensitivity peak for cone vision at the foveola (Sloan, 1939, 1950), even with a coarse testing method, with no tracking of the retinal locus and a stimulus about 1 degree in diameter. Using a smaller stimulus (10 arcmin) and testing with 0.25 degree spacing between test sites, Stiles observed a more distinct sensitivity peak within the central 1 degree radius with a slope of 2.5 dB/degree, surrounded by a plateau (Stiles, 1949). Further testing with clinical perimetry or fundus-controlled perimetry (so-called microperimetry) devices showed that the foveal sensitivity peak sharpens when stimulus size is further reduced to 6 arcmin (Goldman I size), demonstrating that sensitivity testing is prone to summation (Johnson et al., 1981; Khuu and Kalloniatis, 2015; Choi et al., 2016). Clinicians typically use a Goldmann III stimulus (24 arcmin diameter), which activates about 2500 central cones and was shown to be unable to pick up the steep foveolar sensitivity peak (Choi et al., 2016). However, recent microperimetry studies using a Goldman I stimulus reported a slope of about 1.4 dB/degree (Tuten et al., 2012; Khuu and Kalloniatis, 2015; Choi et al., 2016). In our study, we used a stimulus much smaller than Ricco’s area of the foveola. We thus should expect an even steeper slope due to the rapid increase of cone and parasol field size. Indeed, we observed a 5 times steeper slope of about 7 dB/degree within the central 0.4 degree, but with a sensitivity plateau for the central 0.1 degree radius.

An ISETBio model simulating cone activation with different cone spacings was applied to assess the effect of spatial summation. This model predicted a small influence of spacing on sensitivity thresholds due to the size of our stimulus, but did not include any effects due to different light propagation in differently sized cones as proposed by Meadway and Sincich (2018) or increased summation due to an increased ganglion field size.

Furthermore, we found a correlation for the median foveal sensitivity threshold and the participant’s peak cone density. Using a clinical Humphrey field perimeter to assess sensitivity lacking fundus controlled delivery, no correlation between foveolar sensitivity and the minimum cone spacing and therefore maximum cone density with sensitivity in healthy participants was found (Foote et al., 2018; Bensinger et al., 2019). However, assessing sensitivity with a commercial microperimetry system, a significant correlation of sensitivity and density was observed across the central ± 5 degree eccentricity (Agarwal et al., 2015; Supriya et al., 2015; Foote et al., 2019).

### The factor “Outer segment length”

Another factor influencing light sensitivity is the cone OS. Because the photopigment is stored in the OS, its length is directly linked with optical density (Bowmaker et al., 1978; Baylor et al., 1979), and therefore associated with the isomerizations and biochemical transduction cascade. The estimation of the foveal cone’s OS length from a clinical grade OCT device is still part of an ongoing discussion. At first it was reported, that the functional OS is to be found in between the 2nd and 3rd band, since the 2nd band was not the IS/OS junction but the ellipsoid zone of the IS (see also (Lu et al., 2012)) and the 3rd band was not the OS tip, but the contact cylinder (Spaide and Curcio, 2011).

Contradicting these findings it was demonstrated with an AO-OCT that the origin of these two bands are more likely to be the IS/OS junction and OS tips (Jonnal et al., 2014, 2017). Their theory is supported by a computational model simulating the light reflectivity of cones (Meadway and Sincich, 2019). But, there are still uncertainties (see Jonnal et al., 2015; Spaide, 2015), since band 2 and band 3 look very different in AO-OCT and conventional OCT. It was later stated that the thickness of band 3 may be overestimated in a conventional OCT resulting in an overestimation of OS length (Jonnal et al., 2017). Recent findings (Cuenca et al., 2018, 2020; Xie et al., 2018) support the initial report that in conventional OCT band 2 origins from the IS ellipsoids and band 3 from the phagosome zone. That is why we here decided to follow the findings reported in Spaide and Curcio (2011).

Applying these approaches to our OS length estimation we found maximum OS lengths between 28 and 40 µm, which is much smaller compared to other OS length reports for healthy participants from conventional OCT (∼41 µm (N = 43) (Srinivasan et al., 2008); ∼ 47 µm (N = 23) (Wilk et al., 2017b); ∼ 52 µm (N = 97) (Maden et al., 2017); average values), but closer to histological reports (35 µm (Polyak, 1941); >45 µm (Yamada, 1969); 30 µm (Hoang et al., 2002)).

The shortening of the cone OS is accompanied with a decrease of photopigment density over eccentricity (Elsner et al., 1993). In accordance we found that thresholds were well correlated with foveal cone OS length, which was recently observed by Foote et al. (2019), too, but not by Bensinger et al. (2019). Studies examining this relationship of OS length and sensitivity in diseased eyes (Age-related Macular degeneration (Acton et al., 2012; Wu et al., 2014); Glaucoma (Asaoka et al., 2017); retinitis pigmentosa (but IS/OS length) (Mitamura et al., 2009), reported a significant correlation, too.

The ISETBio model confirmed this relationship, based on the assumption that the amount of photopigment per cone is relatively constant (Marcos et al., 1997). Thus, with increasing cone diameter and decreasing outer segment length over eccentricity the resulting optical density decreases.

For the four participants tested herein, we furthermore observed a direct coupling of maximum OS length and the average or peak cone density. Recent publications presented varying results about a correlation of foveal cone density and OS length: While some publications reported a significant correlation for healthy participants (Wilk et al., 2017b; Bensinger et al., 2019; Foote et al., 2019), others found no correlation between maximum cone density and OS lengths (Foote et al., 2018; Allphin et al., 2020).

### Variability of threshold estimates

Taking all these observations together, we developed a model which predicted the shift of a threshold from the average threshold based on the test site’s spacing, OS length and distance from the SC. This model confirmed that all these three factors are important considering sensitivity within the same retina and across different participants. However, this model showed a residual variability of ±0.2 log photons for the expected thresholds which could also not be explained by our ISETBio model for different cone activation patterns or cone classes spectral sensitivity. Also, this variability was more than three times higher than the average variability given by the standard deviation of 0.06 log10 photons we observed during repeated testing (Fig. 3C). For further investigation of this phenomenon, we analyzed replicate testings from a piloting experiment in P2 and extended stimulation of the surrounding retinal area of a cone with conspicuous high thresholds in P3 (Fig. 11). Based on the intra run variability (STD = ± 0.1 log10 photons), threshold differences up to 0.2 log10 photons are not significant. Here, we observed a maximum threshold difference of 0.4 log10 photons between test sites spaced only 1 arcmin apart. These additional data points reveal a very strong dependency between retinal location and small spot sensitivity, presumably caused by different weighting of individual cones. Furthermore, the replicate testing in P2 from another day confirmed that these fluctuations in sensitivity are not only on a small local scale, but also stable for a few days and can be assessed repeatable via AOSLO microstimulation.

**Figure 11:**
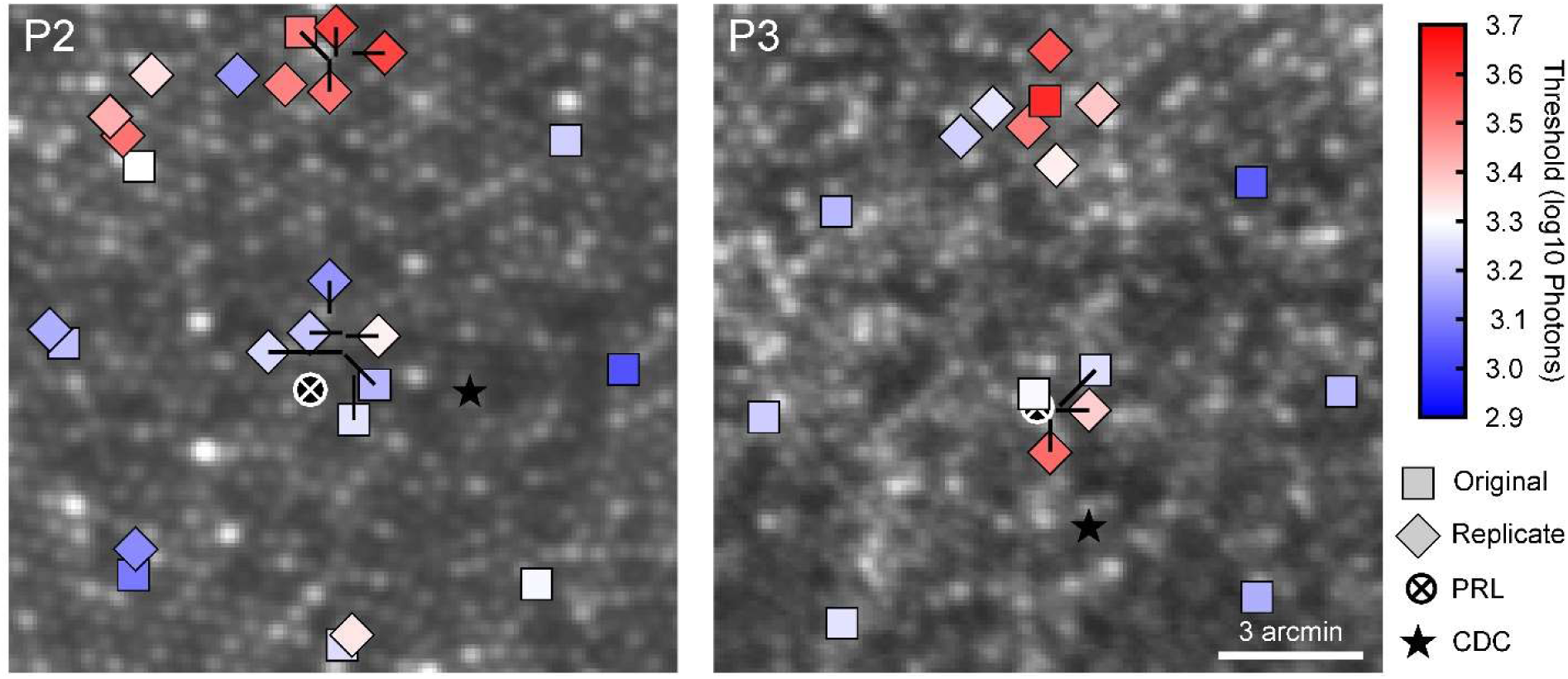
Replicate testing of test sites with conspicuous high thresholds. Squares mark the formerly shown original test sites (Fig. 4), while diamonds mark additionally collected data. The markers match the nominal size of the stimulus. To reduce overlapping, markers were shifted and the tip of a black line indicates the center of the retinal location. For P2 the replicate data points were collected 2 days earlier during a piloting experiment. In P3, the additional locations were tested on the same day, due to the suspiciously high threshold. The PRL is marked with the white circle and crosshair, the cone density centroid by a black star.

This finding of distinct threshold variations between neighboring test sites is consistent with recent work by (Sincich et al., 2016): Using a cone sized stimulus, thresholds changed up to 100 % between neighboring cones (about 0.3 log units). The author’s conclusion was that such unexpected observations might be caused by the individual weighting of each cone.

One possible source of variability could be minimal variations of the photopigment in the individual cones from the same retina (Schnapf et al., 1987, 1990). But this alone cannot explain such high differences between neighboring cones. A more likely explanation could be given by the observation that the individual cone input to ganglion cells varies, due to functional weighting (Chichilnisky and Baylor, 1999; Field et al., 2010; Li et al., 2014). Such functional weighting could happen at the cone-bipolar- or bipolar-ganglion-synapse. So far, *in vitro* studies reported linear and non-linear interaction between cones due to subunits within receptive fields of a ganglion cell (Freeman et al., 2015). Such subunits in the ganglion cell’s field were reported to align directly with the bipolar cells (Liu et al., 2017).

A recent study using AOSLO-based microstimulation found that the actual test site’s cone class composition has a significant influence on sensitivity thresholds (Tuten et al., 2017). The authors reported threshold changes of up to 100 % between test sites with the same cone class composition and the target cone in a surround of a different cone class. They ruled out a simple coupling mechanism through gap junctions (Hsu et al., 2000) and propose a key role of H1 and H2 horizontal cells, mediating an elevated activation level of L-cones due to the 840 nm imaging background thereby inhibiting weak M-cone responses to the stimulus connected to the same horizontal cell (Thoreson and Mangel, 2012). Furthermore, they showed that neighboring S-cones had a suppressing effect on thresholds. The influence of S-cones on this retinal level of modulation during our experiment in the central fovea has to be considered for the following reasons: firstly, histology showed that scattered S-cones can be found within the central 0.15 degrees (Curcio et al., 1991; Bumsted and Hendrickson, 1999), and for individual cases even within the foveola (Ahnelt, 1998). Secondly, our pulsed light source (pulse durations of about 25 psec and a frequency of 100 Mhz) could give rise to activate S-cones via a two photon effect (Palczewska et al., 2014). In the unlikely event of a stimulus being placed centered on an S-cone, the expected threshold would have been elevated by 0.3 log10 photons, based on our stimulation simulation shown in Fig. 10A. Therefore, the conspicuously high threshold found in P3 could be due to an S-cone situated at that location.

On the other hand, sensitivity to small spots could be modulated during post receptor processing in the lateral geniculate nucleus (Jiang et al., 2015; Alitto et al., 2019) or at later cortical stages such as V1 (Gandhi et al., 1999; Smith et al., 2006). The observed high variability between individual test sites might be also an explanation for the reported increase of rerun variability for smaller stimuli (Goldmann V versus Goldmann I) during perimetry, lacking the possibility to stabilize the stimulus on a certain retinal location (Gilpin et al., 1990; Vislisel et al., 2011).

### Estimation of the number of isomerizations at threshold

If we assumed an overall transmission of 41 % for 543 nm light through the ocular media (Boettner and Wolter, 1962), the average threshold would be 2.9 log10 photons at inner segments. Our two-dimensional model of cone capture suggested that the central cones at the target site would have absorbed between 55 photons (7 % of the stimulus light on the retina) and 95 photons (12 %). With an optical density of about 0.5 (Bowmaker and Dartnall, 1980) this corresponds to 27 or 45 isomerizations in either L- or M-cones. The 840 imaging light had a radiant power of about 14.8 log10 photons per second at the cornea (170 µW). With an overall ocular transmission factor of 0.55 [840 nm; (Boettner and Wolter, 1962)] and a field size of 0.85 degree, the estimated photon rate at an individual inner segment was 10.7 log10 photons per second. If a fraction of 33 % was transferred into the outer segments, due to a gaussian absorption model (Macleod et al., 1992) and assuming a cone integration time (cit) of 100 msec (Sperling and Jolliffe, 1965; Krauskopf and Mollon, 1971), we yield roughly 250 R*/cit per L-cone and 22 R*/cit per M-cone. Therefore, increment thresholds found here followed Weber’s law (Reeves et al., 1998) comparing the L-cones isomerizations, but not for M-cones. In other words, we would have expected thresholds to vary by about 1 log unit comparing the isomerizations due to the background, but this was not the case. An explanation could be that even in the central fovea, sensitivity is not transmitted in a 1:1 circuitry but spatially summed across several cones and processed by parasol ganglion cells of the magnocellular pathway (Tuten et al., 2018).

